# Human pluripotent stem cell-derived cardiomyocytes align under cyclic strain when guided by cardiac fibroblasts

**DOI:** 10.1101/2021.02.16.431369

**Authors:** Dylan Mostert, Bart Groenen, Leda Klouda, Robert Passier, Marie-Jose Goumans, Nicholas A. Kurniawan, Carlijn V.C Bouten

**Author notes:** Corresponding author: Carlijn V.C. Bouten, Postal address: Eindhoven University of Technology, Department of Biomedical Engineering, PO Box 513, 5600 MB Eindhoven, The Netherlands. **Competing interest statement:** The authors declare no competing interests.

## Abstract

The human myocardium is a mechanically active tissue typified by anisotropic organization of the resident cells (cardiomyocytes (CMs) and cardiac fibroblasts (cFBs)) and the extracellular matrix (ECM). Upon ischemic injury, the anisotropic tissue is replaced by disorganized scar tissue, eventually resulting in loss of coordinated contraction. Efforts to re-establish tissue anisotropy in the injured myocardium are hampered by a lack of understanding on how CM and/or cFB structural organization is affected by the two major physical cues inherent in the myocardium: ECM organization and cyclic mechanical strain. Herein, we investigate the singular and combined effect of ECM (dis)organization and cyclic strain in a 2D human *in vitro* co-culture model of the myocardial microenvironment. We show that (an)isotropic ECM protein patterning can guide the orientation of CMs and cFBs, both in mono- and co-culture. Subsequent application of uniaxial cyclic strain – mimicking the local anisotropic deformation of beating myocardium – causes no effect when applied parallel to the anisotropic ECM. However, when cultured on isotropic substrates, cFBs, but not CMs, orient away from the direction of cyclic uniaxial strain (strain avoidance). In contrast, CMs show strain avoidance via active remodeling of their sarcomeres only when co-cultured with at least 30% cFBs. Paracrine signaling or N-cadherin-mediated communication between CMs and cFBs were no contributing factors, but our findings suggest that the mechanoresponsive cFBs provide structural guidance for CM orientation and elongation. Our study therefore highlights a synergistic mechanobiological interplay between CMs and cFBs in shaping tissue organization, which is of relevance for regenerating functionally organized myocardium.

## Introduction

The function of the myocardium is highly dependent on its unique structural organization of cells and extracellular matrix (ECM). In the healthy human myocardium, cardiomyocytes (CMs) and quiescent cardiac fibroblasts (cFBs) are arranged as dense, aligned cellular aggregates surrounded by an anisotropic collagen matrix. This spatial arrangement enables electrical and mechanical coupling between CMs and facilitates their synchronous, coordinated contraction [1, 2]. Following ischemic cardiac injury, such as myocardial infarction (MI), billions of CMs die. The ischemia and subsequent inflammatory response at the site of injury initiate the activation and differentiation of cFBs towards myofibroblasts. These cells aid in replacing the damaged tissue with abundant ECM, but also mediate adverse remodeling of the anisotropic myocardium into a disorganized fibrotic scar tissue. In addition, the myofibroblasts secrete profibrotic and hypertrophic cytokines, advancing the progression of adverse remodeling in the continuously beating myocardium, ultimately leading towards heart failure [3 - 5].

The relevance of anisotropic cellular organization for adequate myocardial function has been demonstrated using well-defined *in vitro* systems. For instance, aligned CMs on two-dimensional (2D) substrates demonstrated improved calcium handling and contractile properties compared to randomly oriented CM monolayers [6]. In three-dimensional (3D) cardiac microtissues, aligned CMs and ECM were found to generate a homogeneous and coordinated contraction which was not observed in isotropic tissues [7]. Furthermore, alignment of mesenchymal stem cells, induced by gelatin microgrooves, supported differentiation towards the myocardial lineage [8]. These and other studies suggest that restoring tissue anisotropy may represent a critical step in the functional regeneration of damaged myocardium. Yet, until today cardiac regenerative strategies have largely focused on replenishing CMs [9], while the restoration of the myocardial structural organization is largely overlooked.

*In vivo*, the mechanically active myocardium presents cyclic deformations – due to cardiac beating – as well as structural cues - imposed by collagen fibers - to the CMs and cFBs. Such microenvironmental biophysical cues can induce CM and cFB alignment by mediating the anisotropic organization of their internal structures. *In vitro*, anisotropic structural guidance cues have been imposed using parallel micro-grooves, adhesive protein patterns, or aligned scaffold fibers, demonstrating that both cFBs [10] and CMs [6] can sense this 2D micro-environment and align with the anisotropy presented - a phenomenon termed *contact guidance*. When imposing uniaxial cyclic strain, adherent cells may (re)orient towards the direction (almost) perpendicular to the strain, a phenomenon called *strain avoidance*. Strain avoidance is believed to be cell type dependent, and a minimal strain rate, strain frequency, and cell contractile state is necessary for the behavior to occur [11]. Dynamic reorganization and remodeling of actin stress fibers in combination with a nucleo-cytoskeletal connection is believed to play an important role in a cell’s ability to respond to uniaxial cyclic strain (mechanoresponse) [12, 13]. While cFBs are consistent in their strain avoidance behavior [14], CMs show inconsistent responses, probably related to differences in their contractile state and actin organization [15, 16]. To date, CM mechanoresponse remains largely unexplored, contributing to the controversy about how CMs respond to cyclic strain. Moreover, little is known about the collective organization of CMs and cFBs, and how this is guided or disrupted by structural and mechanical cues in the myocardium [17]. Understanding multi-cell type tissue organization and disorganization is key to not only create accurate *in vitro* models of healthy and diseased myocardial tissue but can also provide deeper insight into the success and failures of *in vivo* myocardial regeneration, including cell-based therapies. To this end, it should become clear how biophysical cues in the myocardium can contribute to the collective organization of CMs and cFBs, and how CMs and cFBs independently contribute to this organization.

In this study, we address this knowledge gap by emulating the fibrous ECM organization and beating of the myocardium in a 2D *in vitro* model of the human myocardial microenvironment. Human fetal epicardial cell derived cFBs and human pluripotent stem cell derived cardiomyocytes (CMs) were cultured on (an)isotropic ECM protein patterns and subsequently subjected to uniaxial cyclic strain. By systematically examining the cellular response of monocultures as well as co-cultures of (varying ratios of) CMs and cFBs to singular and combined environmental cues, this approach provided novel insights into the interplay between the two cell types in regulating organization in physiological and pathological myocardial tissue compositions.

## Results

### ECM patterning guides CM and cFB orientation

Structurally, the myocardium is a fibrous tissue, implying that cardiac cells receive structural cues from contact with ECM fibers. Similar anisotropic topographical or structural cues from the substrate, such as micropatterns, microfabricated grooves, or collagen fibers, have been shown to guide the orientation of CMs and cFBs – a phenomenon termed “contact-guidance” [6, 7, 18, 19]. Here, to mimic the local anisotropic structure of the healthy myocardium and to allow application of mechanical strain, we created fibronectin (FN) adhesive patterns on deformable polydimethylsiloxane (PDMS) membranes using previously established microcontact printing methods (Figure 1) [20]. Two types of patterns were produced: (i) parallel lines with a line width (10 μm), identical to the inter-line spacing (Figure 1A), corresponding to the physiological fiber size of perimysial collagen in the myocardium [21], and (ii) crosshatch patterns (orthogonal linear features of 5 μm linewidth and 10 μm spacing, Figure 1B) to mimic isotropic ECM. The line width for the crosshatch pattern was set to be half that of the parallel line patterns to achieve comparable area density for cell adhesion on the two pattern types. Homogeneously FN-printed PDMS using an unpatterned microstamp served as control substrate (Figure 1C). All FN patterns remained visually intact and well defined throughout the experimental procedures of 48 h of strained or static cell culture (Figure S1).

**Figure 1:**
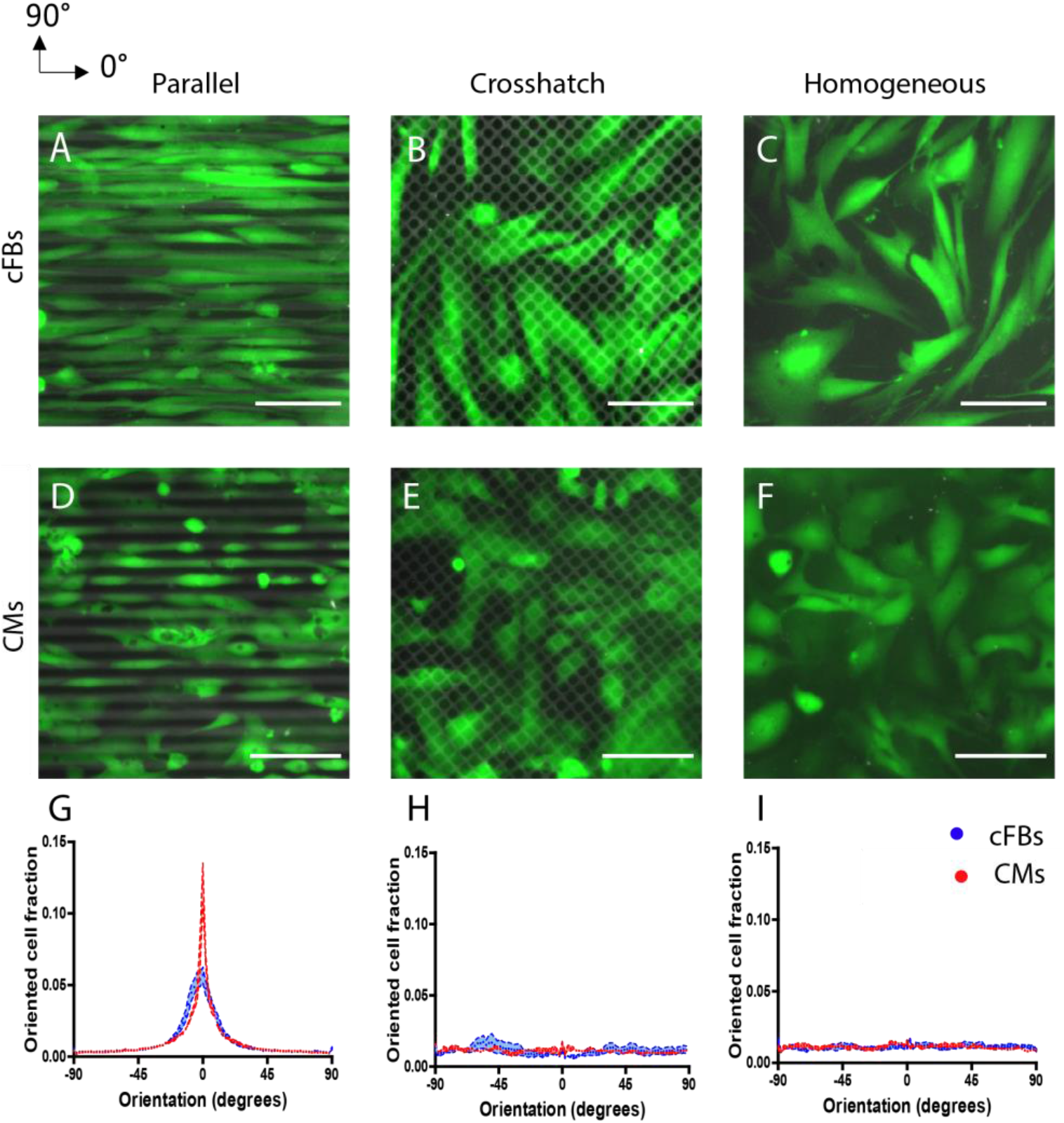
Fibronectin (FN) patterns obtained by micro-contact printing onto deformable PDMS substrates guide cell body organization of cFBs and CMs. A - F) Representative fluorescent images showing FN patterns (TRITC-fibronectin, gray) and cFBs (A-C) and CMs (D-F) (alcein AM, green). G – I) Frequency distribution graphs display the cell orientation with respect to the horizontal direction (0°) for cFBs (blue) and CMs (red). Scale bar indicates 100 μm. Results are expressed as mean ± SEM (n = 9 from 3 independent experiments).

Orientation of cells was evaluated using cytoplasmic calcein AM staining, which circumvents drawbacks with cytoskeleton-based staining due to the substantially different cytoskeletal organizations in CMs and cFBs. Within 24 h of seeding on the printed ECM patterns both cFBs (Figure 1A) and CMs (Figure 1D) showed a clear contact guidance behavior. cFBs were aligned in the direction of the parallel lines and consistently displayed an elongated morphology on all FN patterns with an aspect ratio (AR) 4.5 ± 2.9 on the parallel patterns and 3.5 ± 2.7 on crosshatch patterns. Alignment along the parallel lines was also found for the CMs, although the CMs showed morphological heterogeneity and formed small aggregates on the printed ECM patterns. Such aggregations are often found in contracting CMs cultures and involve cell-cell interactions [22], possibly influencing the amount of CM adhesion to the parallel protein patterns. However, CM density on the substrates did not change during the experiments, suggesting that the vast majority of CMs maintained cell-ECM interactions with the FN patterns. The CMs exhibited an AR of 5.3 ± 1.8 on parallel lines and 2.1 ± 1.0 on crosshatch patterns. The low AR found on the crosshatch protein patterns suggests a relatively immature state of the CMs, derived from stem cells, as opposed to adult CMs *in vivo* [22], which is in line with previous studies using hPSC-CMs [23, 24].

Previously, it has been shown that anisotropic ECM induced the alignment of cFBs [10, 23], hPSC-CMs [25 – 29], and neonatal rat CMs [8, 30]. Consistent with these reports, quantification of cell orientation showed a peak in the direction of the parallel FN lines (0°) and random distribution on crosshatch and homogeneous FN for both cFBs and CMs (Figure 1G-I). Notably, a direct comparison between the two cell types on the same, well-defined ECM patterns, revealed that CMs exhibited more pronounced anisotropic orientation than the cFBs (Figure 1G). This is possibly explained by the smaller cell size of the CMs (110 ± 26 µm longitudinal axis) compared to cFBs (242 ± 59 µm longitudinal axis). Indeed, Buskermolen et al. showed that cell size is an important determinant of cell alignment on microscale ECM patterns and found that the alignment of cardiomyocyte progenitor cells (∼100 μm length) occurred at smaller ECM pattern size compared to that of myofibroblasts (∼200 μm length) [31]. Taken together, these experiments confirmed that ECM patterning is an effective approach to exploit contact guidance to direct the alignment response of both CMs and cFBs.

### cFB cultures, but not CM cultures, show strain avoidance in response to uniaxial cyclic strain

Mechanically, the myocardium and resident cells within experience continuous cyclic mechanical strain due to cardiac beating. Thus, we sought to investigate the orientation response of CMs and cFBs to cyclic strain in the presence of guiding ECM structures mimicked by protein patterns. For several cardiovascular cell types, studies have shown a synergistic effect of structural guidance cues and uniaxial cyclic strain when the cyclic strain direction is presented perpendicular to the anisotropic ECM structures [32, 33]. In contrast, we now asked whether ECM alignment and cyclic strain present competing cues when they are applied in the same direction, similar to the mechanical microenvironment in the myocardium. To this end, we seeded CMs and cFBs on printed ECM patterns that were subsequently subjected to uniaxial cyclic strain at a physiological strain magnitude of 10% and a frequency of 0.5 Hz (Figure 2). In response to such strains, several adherent cell types have been reported to orient perpendicular to the direction of applied cyclic strains – a phenomenon called “strain avoidance” [13, 34 – 36]. On the anisotropic ECM patterns, however, both the CMs and cFBs showed only a slight disruption of cellular alignment without a complete strain avoidance response within the 48 hours of cyclic strain (Figure 2A, D, G). Quantification of the cell orientation demonstrated a slight decrease in the fraction of cells oriented at 0°± 5° (contact-guided response) and a lack of alignment in the direction perpendicular to the strain at 90°± 5° (strain avoidance response) (Figure 2J, M), supporting this dominance of cellular contact guidance over strain avoidance. However, when the starting point was a dis- or nonorganized cellular organization (on crosshatch ECM patterns and homogeneous FN), the cFBs displayed a clear strain avoidance response, as most cells oriented toward 90° (Figure 2B – C, H – I, blue). This is consistent with the reported response of cFBs on homogeneous ECM [14, 37]. In contrast, uniaxial cyclic strain failed to induce orientation of the CMs away from the direction of the applied dynamic strain, even on dis-or nonorganized ECM (Figure 2E – F, H – I, red). This dispersity between cell types is further highlighted by the significant increase in the oriented cell fractions at 90°± 5° for cFBs and the absence of (re)orientation for the CMs (Figure 2N, O). These findings suggest that cFBs and CMs have different intrinsic abilities to sense and respond to cyclic strain.

**Figure 2:**
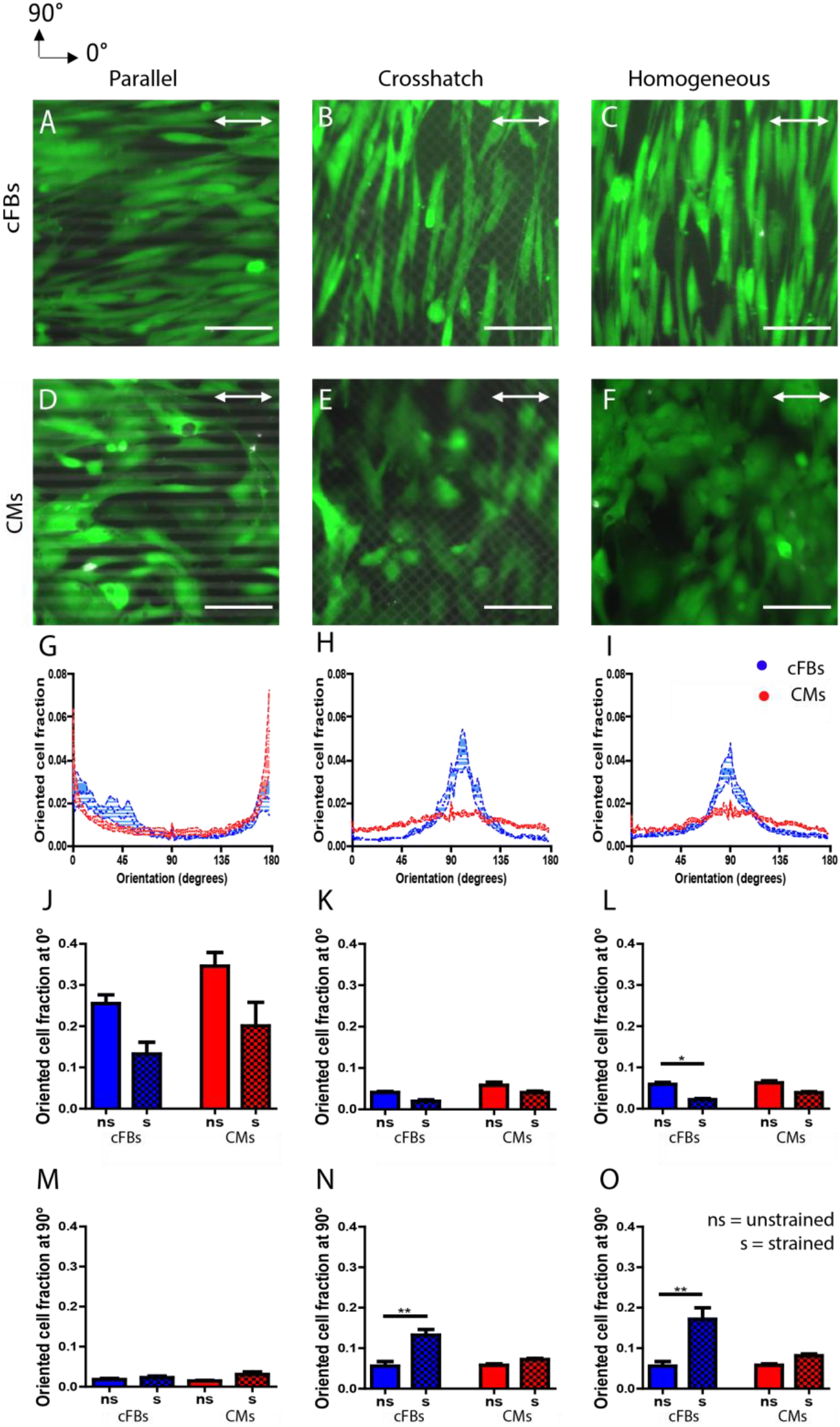
CM and cFB orientation is dominated by contact guidance on anisotropic patterns, while for cFBs strain-avoidance is found on disorganized and homogeneous FN patterns. A - F) Representative fluorescent images showing FN patterns (TRITC-fibronectin, gray) and cFBs (A-C) and CMs (D-F) (Calcein AM, green) after 48 hours of dynamic culture. The direction of the uniaxial cyclic strain is indicated by the white arrow. G - I) Frequency distribution graphs comparing the response of cFBs (blue) and CMs (red) display a clear peak at 90° for cFBs on crosshatch and homogeneous ECM while this is not observed for CMs. J - O) Oriented cell fractions at 0° ± 5° and 90° ± 5°, showing significant reorganization of the cFBs (blue) upon administration of uniaxial cyclic strain whereas this is not observed for CMs (red). ns = non-strained, s=strained. Scale bar indicates 100 μm. Results are expressed as mean ± SEM (n = 9 from 3 independent experiments); * = P < 0.05; ** = P < 0.01.

### Cardiac co-cultures of varying cell ratios organize in response to uniaxial cyclic strain

Given the stark contrast between the responses of CMs and cFBs to environmental cues, we next asked how their difference in mechanoresponsiveness would dictate the collective cellular organization in myocardial co-cultures. To assess how co-culture composition influences the overall mechanoresponse, we created co-cultures of CMs and cFBs with either 70:30 (CM-rich) or 30:70 (cFB-rich) cell seeding ratio. We note that impurities in the CM culture and an increase in number of viable cFBs led to an increased number of non-myocyte cells in the co-culture during the experimental procedure, resulting in co-cultures with 55.3 ± 16.5% α-actinin positive cells (indicative for CMs) for the CM-rich co-cultures and 18.7 ± 9.4% for the cFB-rich co-cultures after 48 hours (Figure S2A). In other words, although the cell composition changed throughout the experimental procedure, the number of CMs was always significantly higher in the CM-rich co-culture. Cell viability remained constant between experimental conditions and time points (Figure S2B - E).

Under static culture conditions, both co-cultures demonstrated cellular alignment on the linear ECM patterns and a random cellular orientation on the crosshatch ECM patterns (Figure 3A, C, E, G), similar to the response of monocultures of cFBs and CMs. Notably, the effect of contact-guidance was found to be more pronounced for the cFB-rich co-cultures than for the CM-rich co-cultures (Figure 3I), suggesting that the presence of cFBs or the crosstalk between cFBs and CMs can influence the susceptibility of either cell type to contact-guidance cues.

**Figure 3:**
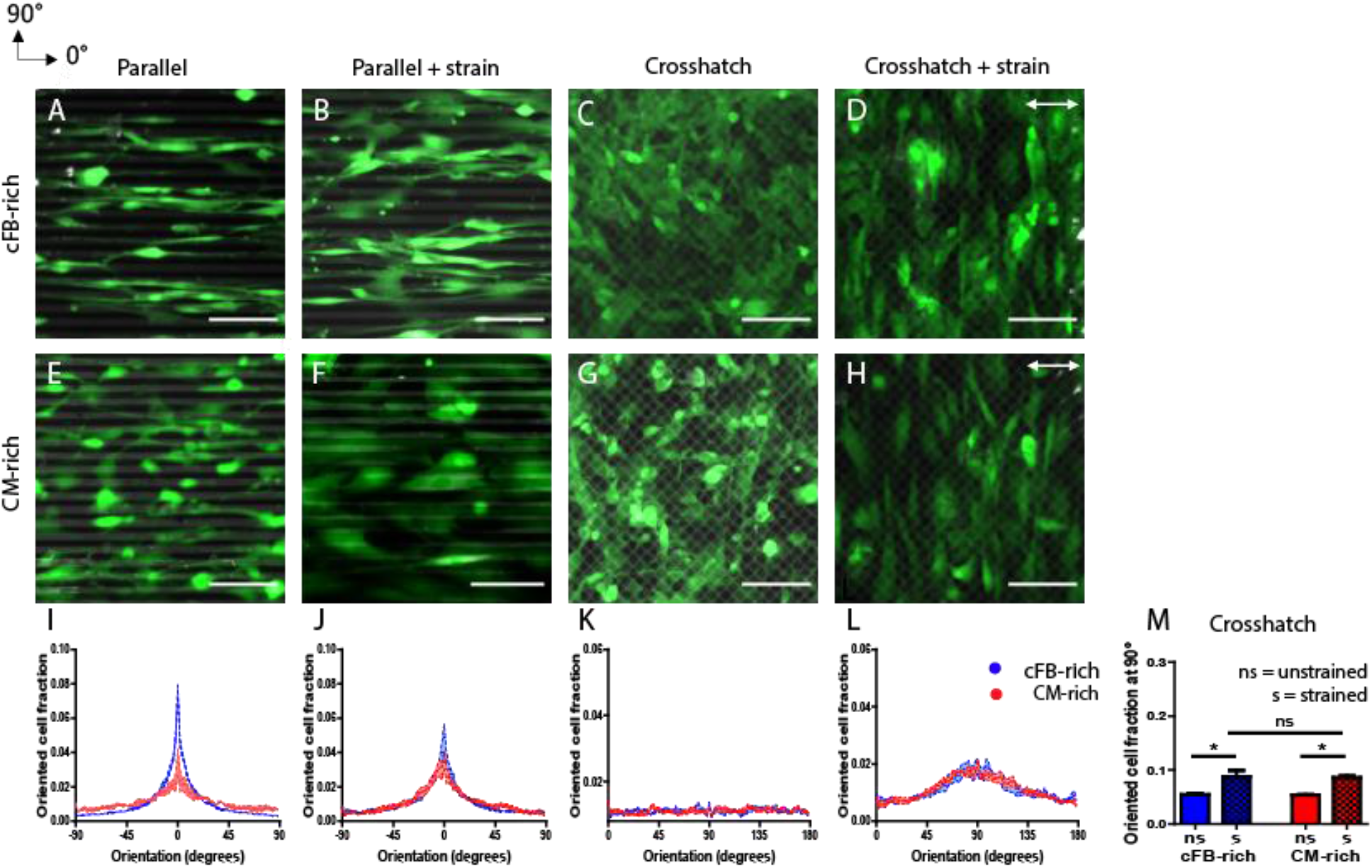
cFB-rich and CM-rich co-cultures show modest strain avoidance behavior in response to uniaxial cyclic strain A-H) Representative fluorescent images (Calcein AM) of the co-cultures after 48 hours of dynamic culture on printed FN patterns. The direction of the uniaxial cyclic strain is depicted with the white arrow. I-L) Frequency distributions histograms comparing the response of cFB-rich co-cultures (blue) and CM-rich co-cultures (red) display the cell orientation corresponding to each co-culture. M) Oriented cell fraction at 90° ± 5° show significant reorganization of the cFB-rich (blue) and CM-rich (red) co-cultures upon administration of uniaxial cyclic strain. Scale bar indicates 100 μm. Ns = non-strained, s = strained. Scale bar indicates 100 μm. Results are expressed as mean ± SEM (n = 9 from 3 independent experiments).

To address how the mechanical environment in the myocardial niche, where ECM guidance and cyclic strain together influence the organization of cardiac cells, affects the organization of cardiac co-cultures, we studied the combinatory effect of uniaxial cyclic strain and (an)isotropic ECM on cFB-rich and CM-rich co-cultures. When uniaxial cyclic strain was applied in the same direction as the anisotropic ECM, both co-cultures lacked a distinct strain avoidance behavior (Figure 3B, F), even though a small but significant increase was found in the fraction of cells aligned at 90° for the cFB-rich co-culture (Figure S3A, C). These results show that contact-guided cellular organization of cardiac co-cultures cannot be easily disrupted by cyclic strain induced strain avoidance, at least with the pattern dimensions and strain protocols used in this study.

Next, we analyzed whether cyclic strain could overpower the disorder that results from disorganized ECM, the cardiac co-cultures were strained after culture on crosshatch patterns. Both the CM-rich and cFB-rich co-cultures showed anisotropic organization of the cellular monolayers after 48 hours of dynamic culture (Figure 3D, H, L), although no significant difference was observed between the oriented cell fraction at 90° of the CM-rich and of the cFB-rich co-cultures (Figure 3M). This was unexpected given that CM monocultures did not show such strain avoidance response. Together, these findings suggest that cFBs guide the orientation of CMs when they align in response to uniaxial cyclic strain, either by direct cell-cell contact or via paracrine signaling. Moreover, these results suggest that uniaxial cyclic strain can be used to induce linear organization of cardiac cells on crosshatch ECM patterns.

To assess the specific contribution of the CMs to the collective reorientation response of the co-cultures, we used a double fluorescent hPSC reporter of mRubyIIACTN2 and GFP-NKX2.5 (DRRAGN) [38]. This cell line demonstrates identical behavior as our earlier findings above, both as monoculture and in co-culture with cFBs (Figure S4). To assess the amount of cFBs necessary to induce the strain-induced alignment of CMs, we used this reporter cell line to systematically create co-cultures with CM-cFB ratios of 20:80, 50:50, 80:20 and subject them to uniaxial cyclic strain for 48 hours on homogenous FN. Qualitative assessment of α-actinin expression by immunofluorescent staining demonstrated aligned sarcomere structures in the direction almost perpendicular to the strain in the 20:80 and 50:50 co-cultures (Figure 4N, S), similar to what was observed in the 70:30 and 30:70 co-cultures. Surprisingly, both CM and cFBs did not reorient in the 80:20 co-culture after strain application (Figure 4U–X). Quantification of sarcomere orientation, which can be linked to the direction of CM contraction, indeed showed a distinct preferred sarcomere orientation in the 90° direction in contrast to unstrained controls (Figure S5G). Besides, as opposed to the more rounded morphologies in CM monocultures and the 80:20 co-culture (Fig 4 D, H, L), CMs demonstrated elongated morphologies and sarcomere alignment both under dynamic and static conditions when enough cFBs were present. Notably, a lack of strain-induced alignment was found for CMs in the 80:20 culture (Figure S5C, F, H), suggesting a threshold of cFBs that is needed in the co-culture to induce strain-induced alignment of CMs. These results suggest a threshold of cFB number (>30%) required for inducing strain avoidance in CMs as well as a more elongated phenotype with aligned sarcomeric structures, indicative of CM maturity [39, 40].

**Figure 4:**
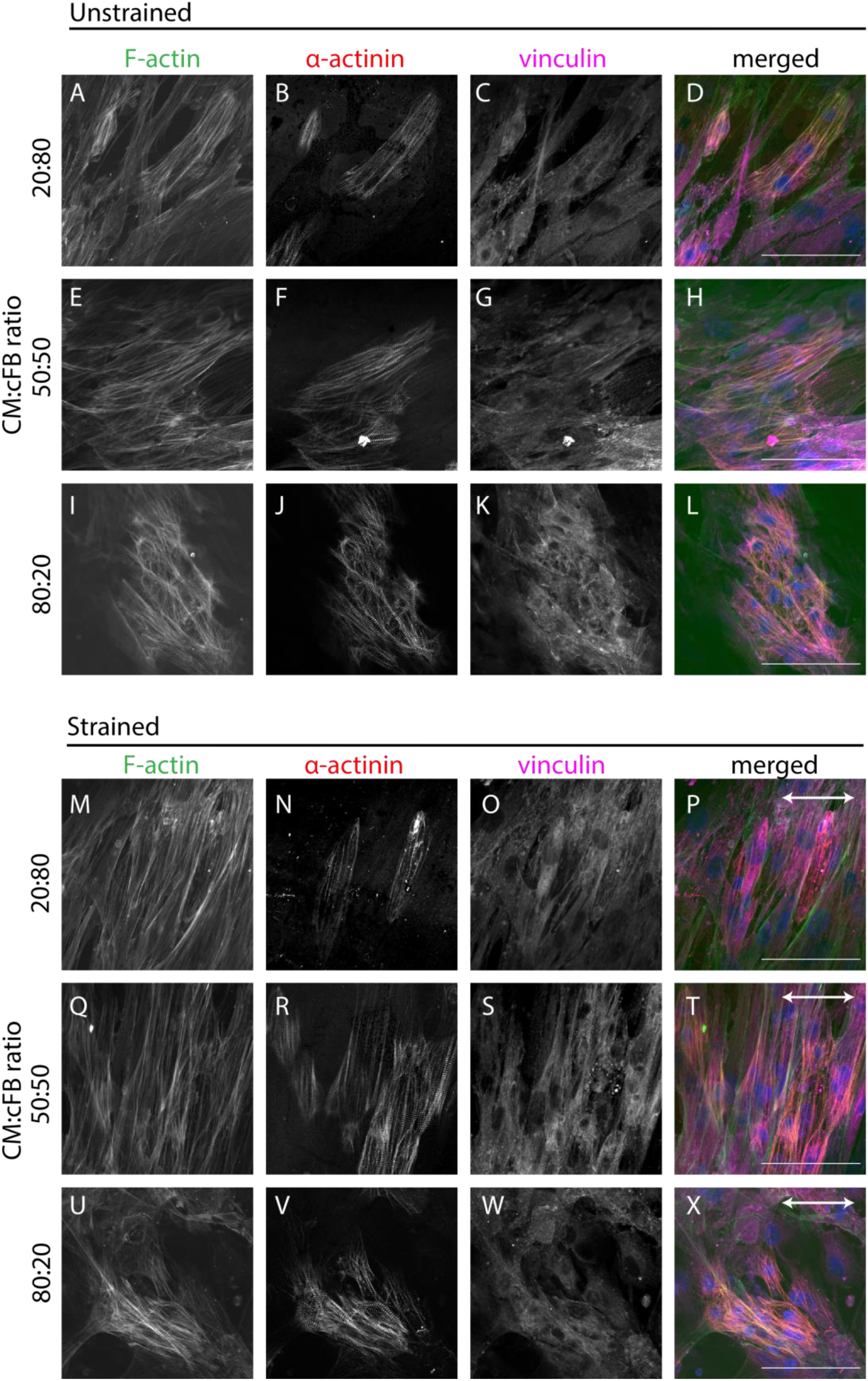
Uniaxial cyclic strain induces sarcomere alignment in CMs when in co-culture with a threshold value of cFBs. A-L) Representative fluorescent images of F-actin (green), α-actinin (red), and vinculin (pink) in co-cultures with varying seeding ratios after 48 hours of static culture on homogeneous FN. M-Z) Representative fluorescent images of the co-cultures after 48 hours of dynamic culture on homogeneous FN. The direction of the uniaxial cyclic strain is depicted with the white arrow. Scale bar indicates 100 μm.

### Strain avoidance of CMs in the presence of cFBs is not mediated by paracrine signaling or N-cadherin

We next asked what mechanisms are underlying the strain avoidance of CMs when in co-culture with a threshold value of cFBs. In particular, we wondered if the observed effect could be explained by paracrine signaling of the cFBs or by direct cell-cell communication via adherens junctional protein N-cadherin. To assess whether the impact of cFBs on CMs alignment could be attributed to paracrine factors, conditioned medium was collected from cFB monocultures after 48 h of uniaxial cyclic strain and added to the CMs before strain application. After 48 h cyclic strain of CMs in the conditioned medium, no significant differences in cell orientation were found in comparison to untreated monocultures of CMs (Figure S6A, B, E, F). Moreover, limited sarcomere formation was observed in the CMs in the conditioned medium, suggesting that the improved maturation state of CMs in co-culture with cFBs is not attributable to paracrine signaling (Figure S6C, D). To investigate whether adherens-junction-mediated cell-cell communications play a role in the strain-induced alignment of CMs in cardiac co-cultures, we analyzed the expression and localization of N-cadherin (Figure S7). While clear N-cadherin expression was observed at the cell membrane between CMs, N-cadherin was absent between cFBs. Moreover, although CMs express cytoplasmic N-cadherin when in direct contact with cFBs, they lack the clear membrane bound N-cadherin expression found between CMs (Figure S7G - K). For Connexin43 (Cx43), the most abundant gap junctional protein in CMs, similar cytoplasmic expression was found as for N-cadherin and no membrane bound Cx43 was observed in between CMs and cFBs (data not shown). These results indicate that the strain-induced alignment of CMs in co-culture with cFBs is not caused by paracrine signaling nor to Cx43 or N-cadherin mediated cell-cell communication between the CMs and cFBs. This is in line with a recent report showing a lack of Cx43 and N-cadherin-mediated cell-cell communication in the co-culture organization of neonatal rat CMs and cFBs [41]. Future experiments using targeted inhibition of Cx43 and N-cadherin in co-cultures under cyclic strain together with transcriptional analysis should be performed to validate these findings.

## Discussion

In our search to develop strategies to regenerate cardiac anisotropy, we aimed to gain fundamental understanding of how biophysical cues from the myocardium contribute to shaping and disrupting anisotropy by organizing the main cell types in the cardiac wall. Specifically, we studied the collective reorganization response of cFBs and CMs, derived from induced pluripotent stem cells, under the influence of cyclic strain and ECM protein patterns to model both the cellular composition of myocardial tissue in health and disease as well as the dynamic CM microenvironment. Notably, most studies that address the effect of biophysical cues on cardiac cell organization have focused solely on monocultures [6, 10, 42 – 44], while crosstalk between cFBs and CMs is found to be critical for myocardial function [45, 46]. To this end, we used a 2D *in vitro* approach of micropatterned ECM protein patterns on deformable substrates, which allowed controlled presentation of structural cues and mechanical strains independently and simultaneously to monocultures and co-cultures of cFBs and CMs. We show that CMs in co-culture with ≥ 30% cFBs exhibit a collective strain avoidance response concurrent with sarcomere alignment in the same direction, which was not found for the CM monocultures. Moreover, we observed that uniaxial cyclic strain could not induce alignment of CMs via solely paracrine signaling of the cFBs or when the cFBs lacked a strain avoidance response. Given this evidence, we hypothesize that uniaxial cyclic strain promotes strain avoidance in cFBs, which in turn act as guidance cues for hPSC-CMs (Figure 5).

**Figure 5:**
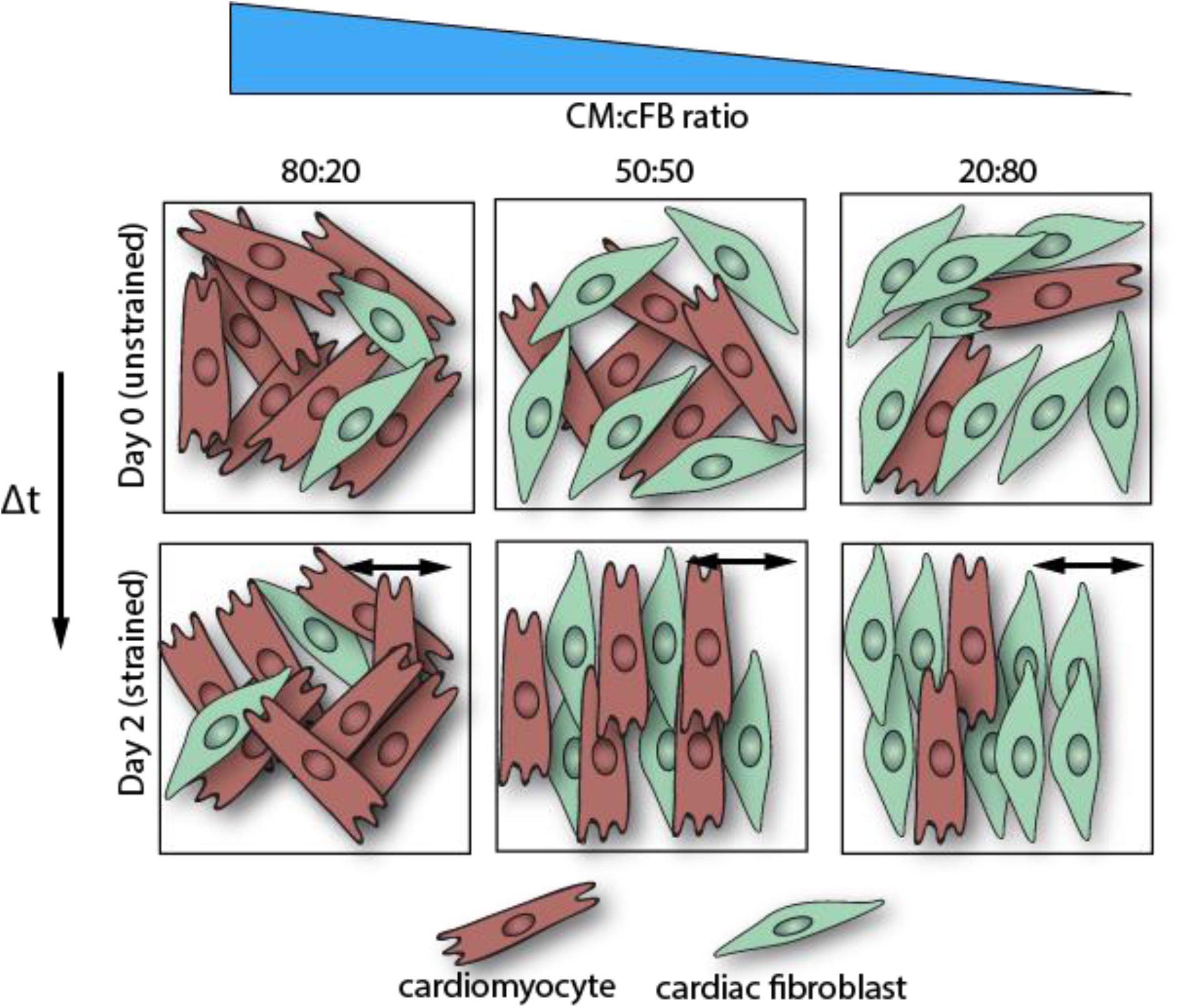
Conceptual schematic representation of the changes in cFB orientation upon application of 48 hours of uniaxial cyclic strain and the proposed link to CM alignment. Top row: co-cultures of various seeding densities show random organization under static culture conditions. Bottom row: Upon cyclic strain application, strain-avoidance behavior is found for cFBs, which in turn serve as guidance cues for hPSC-CM orientation. We propose the formation of anisotropic guidance cues, resulting from the strain-avoidance response of cFBs, which steer hPSC-CM alignment and sarcomere organization in the direction almost perpendicular to the applied cyclic strain.

Analysis of CMs and cFBs in monoculture and in co-cultures, seeded in a 70/30 or 30/70 (CM/cFB) ratio, on ECM patterns under static conditions revealed that structural cues can effectively guide cellular orientation, resulting in anisotropic cell organization on parallel ECM patterns. Since myocardial tissue mainly consists of parallel collagen I fibers, ECM organization could be a key determinant of cFB and CM organization *in vivo* [47]. Strikingly, distinct responses were observed for cFBs and CMs when ECM patterns and uniaxial cyclic strain were presented simultaneously. Whereas cFBs on homogeneous and crosshatch ECM patterns showed a clear strain avoidance response, we found that monocultures of CMs did not show strain avoidance. This is in contrast with Qi et al. and Kreutzer et al., who observed stain avoidance for hPSC-CMs [43, 44]. These inconsistencies might be attributed to the small FAs found in monocultures of CMs or lower expression levels of fibronectin binding integrins [17], which could hamper proper transduction of mechanical forces from the microenvironment to the CMs. Moreover, in the present study we used lower strain rates and straining time than those used in earlier reports, which might affect the strain avoidance response. Also, differences in CM maturity might contribute to the lack of strain-induced alignment since differentiation state is found to influence strain avoidance behavior in cardiomyocyte progenitor cells [36]. Despite the lack of response by CMs, the clear induction of anisotropic organization on the crosshatch patterns for cFBs suggests that uniaxial cyclic strain can be exploited to restore the anisotropic architecture of the myocardium if cFBs dominate the injured area of the myocardium.

Whereas, to our knowledge, no literature exists for human tissues or models on how the amount of cFBs modulates strain avoidance of CMs, a recent study by Tran et al. demonstrated a cell density dependent response to cyclic strain for co-cultures of varying ratios of neonatal rat CMs and cFBs [41]. In contrast to hPSC-derived CMs, monocultures of neonatal rat CMs oriented along the direction of cyclic strain and increasing the amount of cFBs in the co-culture resulted in orientation away from the direction of the applied cyclic strain. However, a higher number of cFBs was needed to induce strain avoidance in neonatal rat CMs as opposed to in co-cultures of human cFBs and CMs derived from pluripotent stem cells, possibly due to differences in maturity and contractile state between the two types of CMs [41]. Moreover, several studies have investigated the contribution of the amount of cFBs on CM alignment and maturity. For CM maturity, an optimum in contractility was found in co-cultures containing 10-30% cFBs, which could not be achieved when dermal fibroblasts were used instead of cFBs [48]. Also, for neonatal rat CMs in cardiac microtissues, a cFB density of 55% or more weakened CM function, demonstrated by a significant decrease in contractile force [49]. Interestingly, Rupert et al. showed that 15% of cFBs in co-culture with hPSC-CMs in engineered cardiac tissues resulted in less CMs alignment as opposed to 5% of cFBs [50], seemingly in contrast to the observations in the present study. However, the experiments performed by Rupert et al. were conducted in engineered cardiac tissues where cFBs adopt a more quiescent phenotype as opposed to the contractile phenotype on stiff 2D substrates [50], suggesting that the phenotype of cFB may play a role in aligning CMs. These findings demonstrate that a low percentage of cFBs improves CM maturation and tissue contractility. Yet, the diseased myocardial microenvironment consists of mainly contractile cFBs, suggesting that the mechanoresponsiveness of cFBs can be critical in determining CM and overall tissue organization.

Our findings indicate that CMs and cFBs, the main cellular components of the myocardium, differentially respond to structural and mechanical cues that typify the myocardial tissue. Importantly, an interplay between the two cell types is necessary for the overall strain-induced response that leads to anisotropic cell organization as is expected in healthy, contractile myocardium. Therefore, our results propose an intriguing emergence of guidance cues for CMs that are mediated by the mechanoresponsiveness of cFBs, Whereas cell orientation can be dynamically induced by topographies and structural guidance cues in *in vitro* culture using light-responsive synthetic materials [51 – 53],we present a similar emergence arising from cues that are inherent in the *in vivo* CM microenvironment – mechanical strain and cFBs. Further investigation should address if removal of cyclic strain can cease the presentation of the cFB-mediated guidance cues, which would suggest dynamic control over the presentation of these guidance cues.

Collectively, our data propose the importance of the strain avoidance response of cFBs, a cell type often overlooked in cardiac regenerative strategies, in determining myocardial architecture and function. Future studies should indicate how the mechanoresponsiveness of cFBs can be exploited in 3D and in *in vivo* heart models as a strategy to regain cardiac anisotropy upon injury. These results not only shed new light on the role of cFBs in structural remodeling upon myocardial damage, but also point at a possible role of cFBs in regaining myocardial anisotropy in the damaged heart. To apply the latter option, computational models should be employed that explore how mechanical strains present in the infarcted myocardial tissue can be used and manipulated to organize the CMs and cFBs along the direction of contraction[54, 55]. Moreover, by exploiting the strain avoidance response of cFBs, the formation of anisotropic structure in engineered tissue constructs can be promoted, aiding the design of strategies for structural organization of the myocardium.

## Materials and Methods

### Fabrication of micro-patterned deformable substrates

Fibronectin (FN) adhesive patterns were transferred onto deformable PDMS membranes using previously established protocols for microcontact printing [20, 56]. In short, a parallel line pattern (10 μm width, 10 μm gap) and a crosshatch pattern (perpendicular crossing with 5 μm wide lines with 10 μm spacing) were generated on a silicon master wafer by deep reactive-ion etching from a chromium photomask [57]. Microstamps were obtained by molding the silicon master with polydimethylsiloxane (PDMS, Sylgard 184; Dow Corning, Michigan, United States) and curing agent (10:1) and curing the constructs for 20 minutes at 120 °C. Cured PDMS microstamps with the desired features were then peeled off from the master and cleaned and dried with 70% ethanol and compressed air, respectively. In order to print the protein patterns, the microstamps were inked for 1 hour at room temperature with 25 μg/mL rhodamine FN solution (Cytoskeleton Inc., Colorado, United States). Deformable silicone Flexwell membranes (Flexcell inc., Burlington, United States), supported by an underlying printing mold, served as substrates for the microcontact printing. These substrates were first oxidized in an UV/ozone cleaner (PDS UV-ozone cleaner; Novascan, Iowa, United States) for 8 minutes, after which the FN-coated stamps (first dried under compressed air) were gently deposited on the substrates and incubated for 15 minutes at room temperature. Uncoated regions were blocked by immersing the substrates for 5 minutes in 1% Pluronic F-127 (Sigma-Aldrich, Missouri, United States) in PBS. Finally, the membranes were washed three times with phosphate-buffered saline (PBS, Sigma-Aldrich) and stored in PBS at 4°C until further use. A microstamp without any features was used to print homogeneous FN as a control substrate.

### Human induced pluripotent stem cell derived cardiomyocyte culture

Human induced pluripotent stem cell derived cardiomyocytes (hiPSC-CMs) were differentiated according to protocol described before [58] and kindly provided by Prof. E. van Rooij (Hubrechts Institute, Utrecht, the Netherlands). hiPSC-CMs were maintained in RPMI 1640 with HEPES and Glutamax (Life Tech, Bleiswijk, the Netherlands) with 2% B27 with insulin (Life Tech) and 1% penicillin/streptomycin (P/S, Sigma-Aldrich). Six days before mechanical stimulation, hiPSC-CMs were plated on the micropatterned deformable substrates in order to allow cryopreservation recovery and spontaneous beating retrieval. Cells were seeded at 50.000 cells/cm^2^.

### Human pluripotent stem cell derived cardiomyocyte culture (DRRAGN)

The DRRAGN hPSC (derived from human embryonic stem cell line HES3) line 3F4, modified with a double reporter of GFP-NKX2.5 and mRubyIIACTN2 was kindly provided by prof. R. Passier (University of Twente, the Netherlands). The hPSCs were cultured on vitronectin-coated culture plates and maintained by refreshing Essential 8 (E8) medium (Thermo Fisher, Waltham, Massachusetts, United States) daily and passaging the cells at around 150k cells/cm^2^ each Monday and Thursday. The hPSCs were differentiated into cardiomyocytes (hPSC-CM) according to protocol described before [59]. On day 16 of differentiation, the cells were dissociated and cryopreserved in 50% knock-out serum, 40% BPEL medium, 10% DMSO and rock inhibitor (Y27632, 1:200). The hPSC-CMs were thawed five days before the start of each experiment.

### Epicardial-derived cardiac fibroblast culture

Human epicardial-derived cardiac fibroblasts (cFBs) were kindly provided by Prof. M.J.Th.H. Goumans (Leiden University Medical Centre, the Netherlands). cFBs were isolated from human fetal cardiac tissue collected with informed consent and anonymously from elective abortion material of fetuses with a gestational age between 10 and 20 weeks. This research was carried out according to the official guidelines of the Leiden University Medical Center and approved by the local Medical Ethics Committee (number P08.087). cFBs were cultured in high-glucose Dulbecco’s modified Eagle’s medium (Invitrogen, Breda, the Netherlands) supplemented with 10% fetal bovine serum (Greiner Bio-one) and 1% P/S (fibroblast culture medium). The cFBs were cultured in flasks coated with 0.1% gelatin from porcine skin in PBS (Sigma-Aldrich) and passaged at 80% confluency. The passage number of the cFBs during mechanical stimulation was between 4 and 8.

### Cardiac co-cultures

In order to analyze the contribution of cFBs to the alignment response of CMs, co-cultures were created with varying cell seeding densities. Either hiPSC-CMs or DRRAGN CMs were dissociated with 10x TrypLE (Gibco, Eindhoven, the Netherlands) and seeded in maturation medium (TDI) 48 hours prior to mechanical stimulation (day-2). After 24 hours (day-1) the cFBs were dissociated using 0,05% Trypsin/EDTA (Gibco, Eindhoven, the Netherlands) and added to hPSC-CMs in co-culture medium containing 25% fibroblast culture medium and 75% TDI medium. 150 μL of cell suspension was seeded only on top of the coated region of the flexible PDMS substrates and incubated at 37°C overnight to allow local cell adherence before extra co-culture medium was added. The co-cultures were maintained in hPSC-CM medium and incubated at 37°C and 5% CO_2_ until analysis. Monocultures of hPSC-CMs and cFBs were seeded onto the same substrates and mechanically stimulated as control.

### Cyclic straining of mono and co-cultures

The FX-5000™ Tension System (Flexcell inc., Burlington, United States) was used to provide controlled uniaxial cyclic strain to the cardiac cells cultured on the flexible membranes by means of a regulated vacuum pressure that deforms the membrane. The cardiac cultures were seeded one or two days prior to straining to allow for cell adhesion and cellular organization through the printed fibronectin patterns (d_-1_) and were strained for 48 hours thereafter. Strains of 10% (0.5 Hz, sine wave) were applied to the cardiac cultures, increasing stepwise from day 0 to condition the cells, and reaching 10% strain after 8 hours. On day 0 (before the onset of strain) and after 48 hours (day 2) the samples were collected. Unstrained cardiac mono and co-cultures were used as control. The cyclic strain applied to the membranes in 2D was validated using image analysis. First, graphite powder was sprayed on top of membranes that were excluded from cell culture and several stretching cycles were recorded by a camera and the applied cyclic strain was measured by the displacement of the graphite pattern with ImageJ.

### Cardiac fibroblast conditioned medium

cFB conditioned medium was obtained from cFBs after 48 hours of mechanical uniaxial cyclic strain to allow secreting of paracrine factors. 50.000 cFBs were plated on homogenous substrates 24 hours before mechanical stimulation. 3mL cFB culture medium (as described in 2.3) was added to every well and the cFBs were strained (as described in 2.6) for 48 hours. After mechanical stimulation the conditioned medium was retrieved from the cFBs and filtered using a 0,2 μm syringe filter to remove cells and other large particles. The medium was frozen immediately and thawed just before it was mixed 1:1 with fresh TDI medium and added to monocultures of 100.000 hPSC-CMs, which were mechanically loaded or statically cultured as control.

### Quantification of cell and sarcomere orientation

The orientation response of the cardiac cells was determined from triplicates of three independent experiments (n = 9). Cells were incubated with 1 μg/mL calcein AM (Sigma-Aldrich) for 20 min and the orientation of their long axis was analyzed after medium renewal with an inverted microscope (Zeiss Axiovert 200m equipped with an AxioCam HR camera; Zeiss, Sliedrecht, the Netherlands). Only the central parts of the wells, where the strain field was homogeneous as determined using our in-house calibrations, were considered (1 cm^2^, 50,000 cells). Cell orientation was quantified with ImageJ, using the Fiji plug-in “directionality”, based on Fourier spectrum analysis (Figure S8).

### Viability assay

To verify the coexistence of hiPSC-CMs and cFBs in the co-cultures, cell viability was assessed using calcein AM and propidium iodide (Invitrogen) one day after seeding (day 0) and 48 hours after that (day 2). In short, the co-cultures were carefully washed with PBS and incubated with 1 μg/mL calcein AM and 750 nM propidium iodide for 20 minutes at 37° C protected from light. After incubation, the cultures were washed with PBS directly imaged with an inverted fluorescence microscope (Zeiss Axiovert 200m). Images were taken at three random representative spots per condition. Three individual experiments were performed with 2 – 3 replicates (n = 2 – 3) and co-cultures treated with ethanol were used as negative control.

### Immunofluorescence staining

The cardiac cells cultured on the Bioflex plates were washed with PBS trice before and after fixation, and fixed in 3.7% formaldehyde (Merck, Darmstadt, Germany) for 15 min. The flexible membranes were cut out of the Bioflex plates, and subsequently cut in quarters to allow multiple analysis per well. The cells were permeabilized with 0,5% Triton-X-100 (Merck) in PBS for 10 minutes and blocked for non-specific antibody binding with 10% horse serum (Sigma-Aldrich) in PBS for 40 minutes. The cells were incubated overnight with the primary antibodies at 4°C. The cells were washed 6 times with PBS for 5 minutes before incubating with the secondary antibodies in PBS and phalloidin for 1.5 hours. All used primary and secondary antibodies and dyes are listed in Table S1. The cells were washed 2 times with PBS before incubating with DAPI in PBS for 5 minutes to visualize the cell nuclei. Finally, cells were washed 4 times with PBS for 5 minutes, and thereafter the membranes were mounted to glass slides with Mowiol (Sigma-Aldrich) and stored protected from light at 4°C. All immunofluorescent samples were analyzed with a confocal fluorescence microscope (Leica SP5X) using the 10x and 20x objectives.

### Statistical analysis

Cell orientation data are presented as mean ± standard error of the mean (SEM). Statistical analysis was performed using Wilcoxon matched-pairs signed rank test for non-parametric two group comparison. Significance was assumed when p < 0.05. Graphics and statistical analyses were performed with Graphpad Prism software (version 5.04).

## Acknowledgements

This research was financially supported by the Gravitation Program “Materials Driven Regeneration”, funded by the Netherlands Organization for Scientific Research (024.003.013). dr. N. A. Kurniawan acknowledges financial support from the European Research Council (851960).

## Supplementary Information for

**Figure S1:**
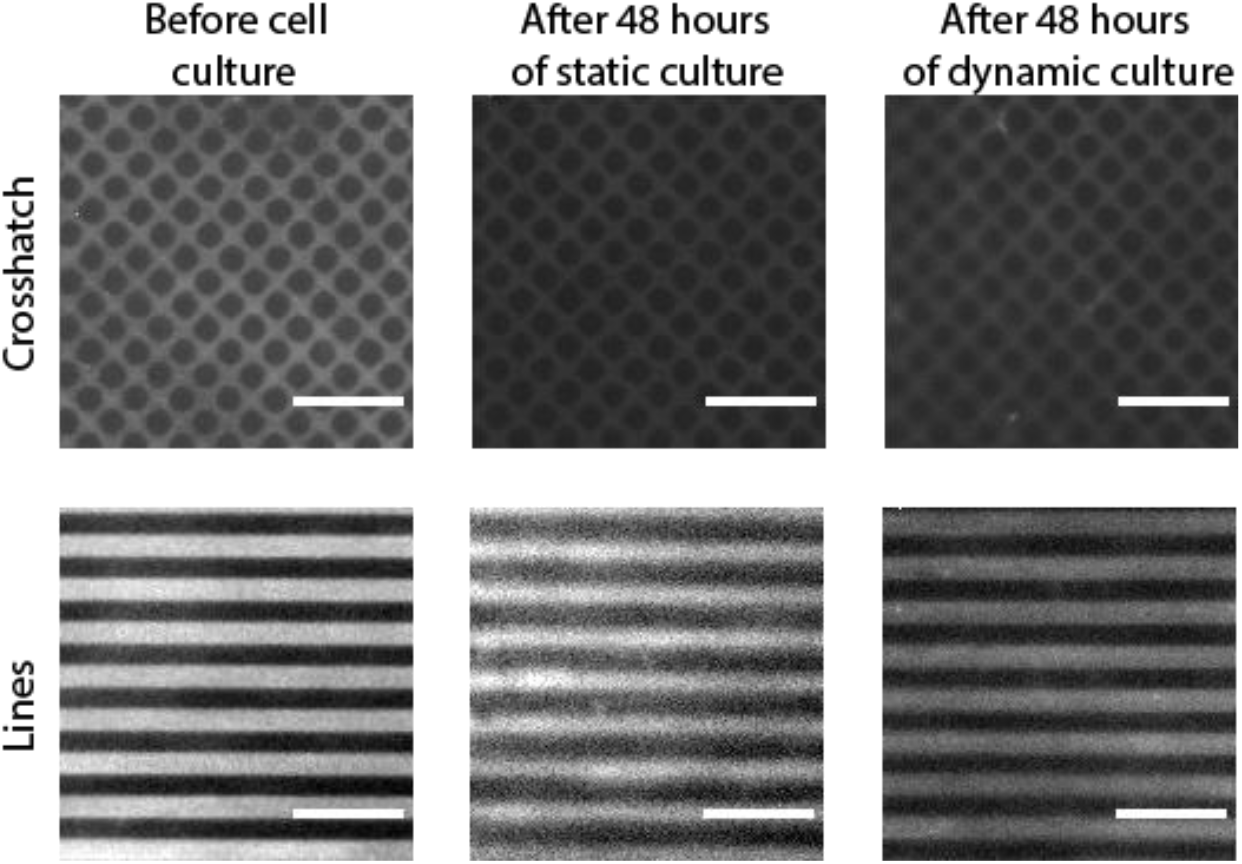
Fibronectin (FN) protein patterning through micro-contact printing demonstrates well-defined and visually intact FN patterns throughout the experimental procedure of 48 hours of static or dynamic culture.

**Figure S2:**
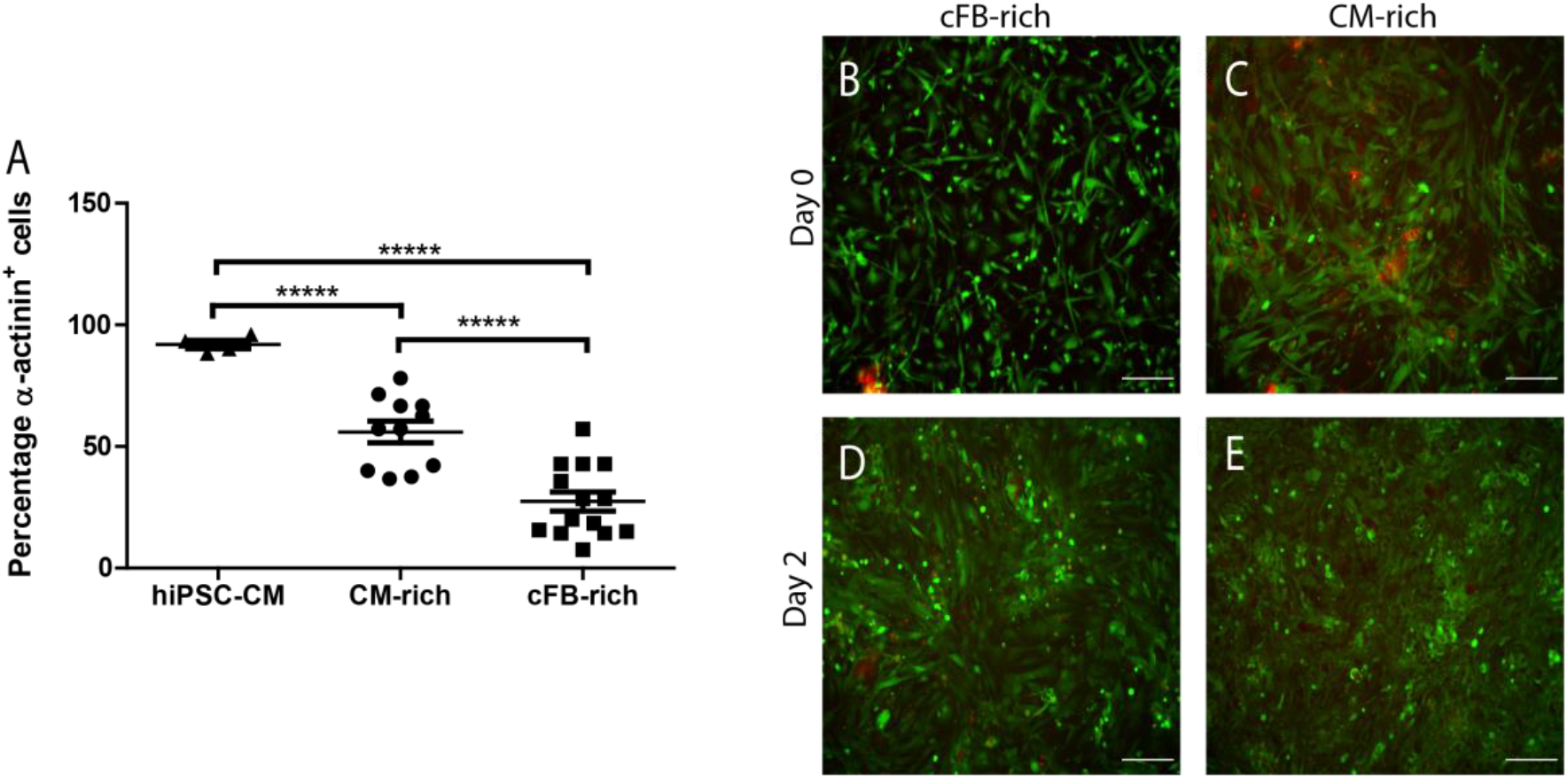
Cellular distribution and viability of cardiac co-cultures over time. A) The percentage of α-actinin positive cells indicates a significant higher amount of CMs in the CM-rich co-culture as opposed to the cFB-rich co-culture over time. B – E) Representative fluorescent images of the cardiac co-cultures showing live cells in green (calcein AM) and dead cells in red (propidium iodide) 24 hours after seeding (day 0) and after 48 hours of culture (day 2) on homogeneous fibronectin coating. Scale bar = 200 μm. **** = P < 0.0001, n = 180 cells.

**Figure S3:**
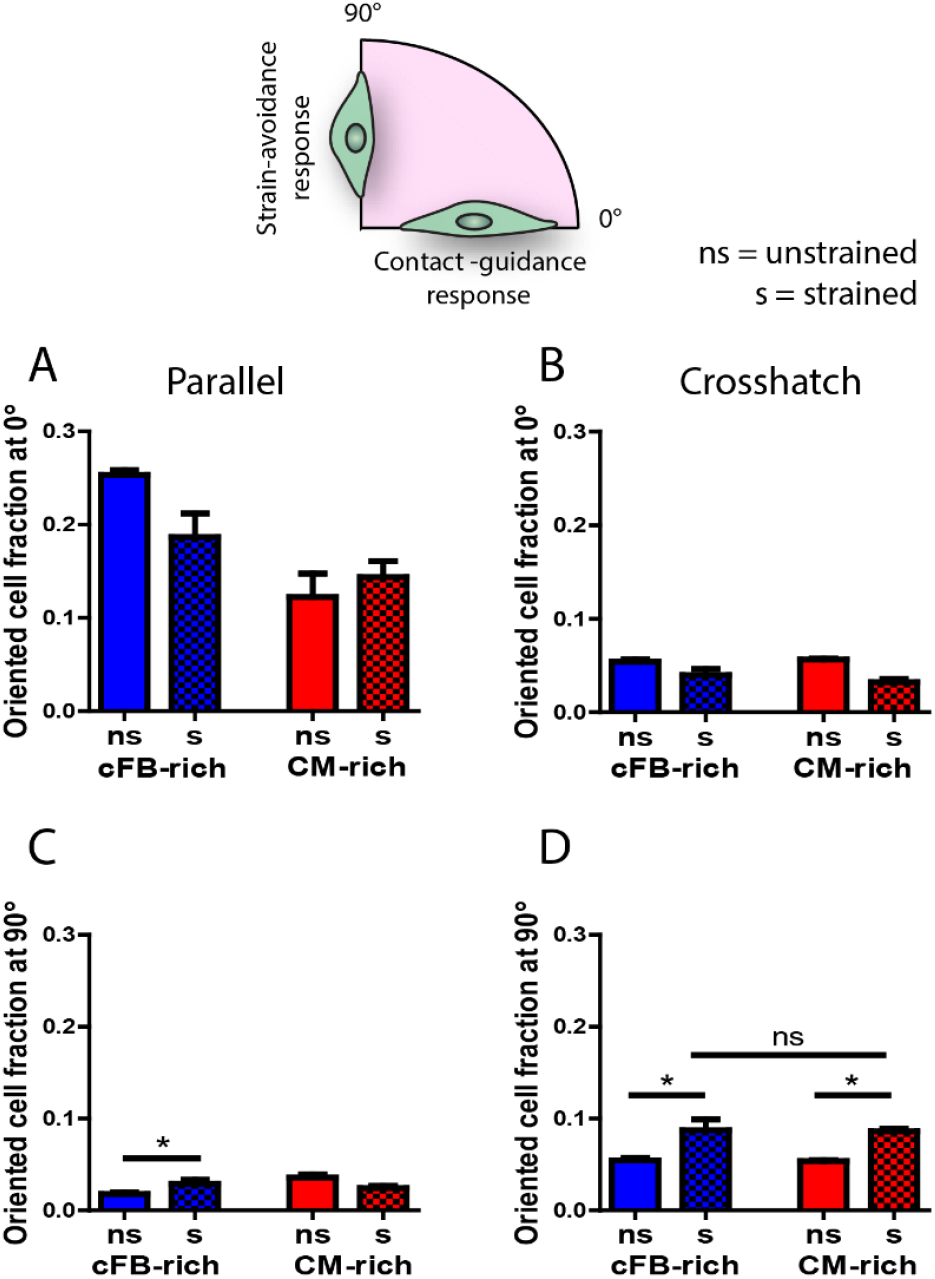
Uniaxial cyclic strain induces the alignment of both CM-rich and cFB-rich co-cultures on crosshatch ECM. A - D) Oriented cell fractions at 0° ± 5° and 90° ± 5° show significant reorganization of the cFB-rich (blue) and CM-rich (red) co-cultures upon administration of uniaxial cyclic strain on the crosshatch patterns. Ns = non-strained, s = strained. Results are expressed as mean ± SEM (n = 9 from 3 independent experiments); * = P < 0.05.

**Figure S4:**
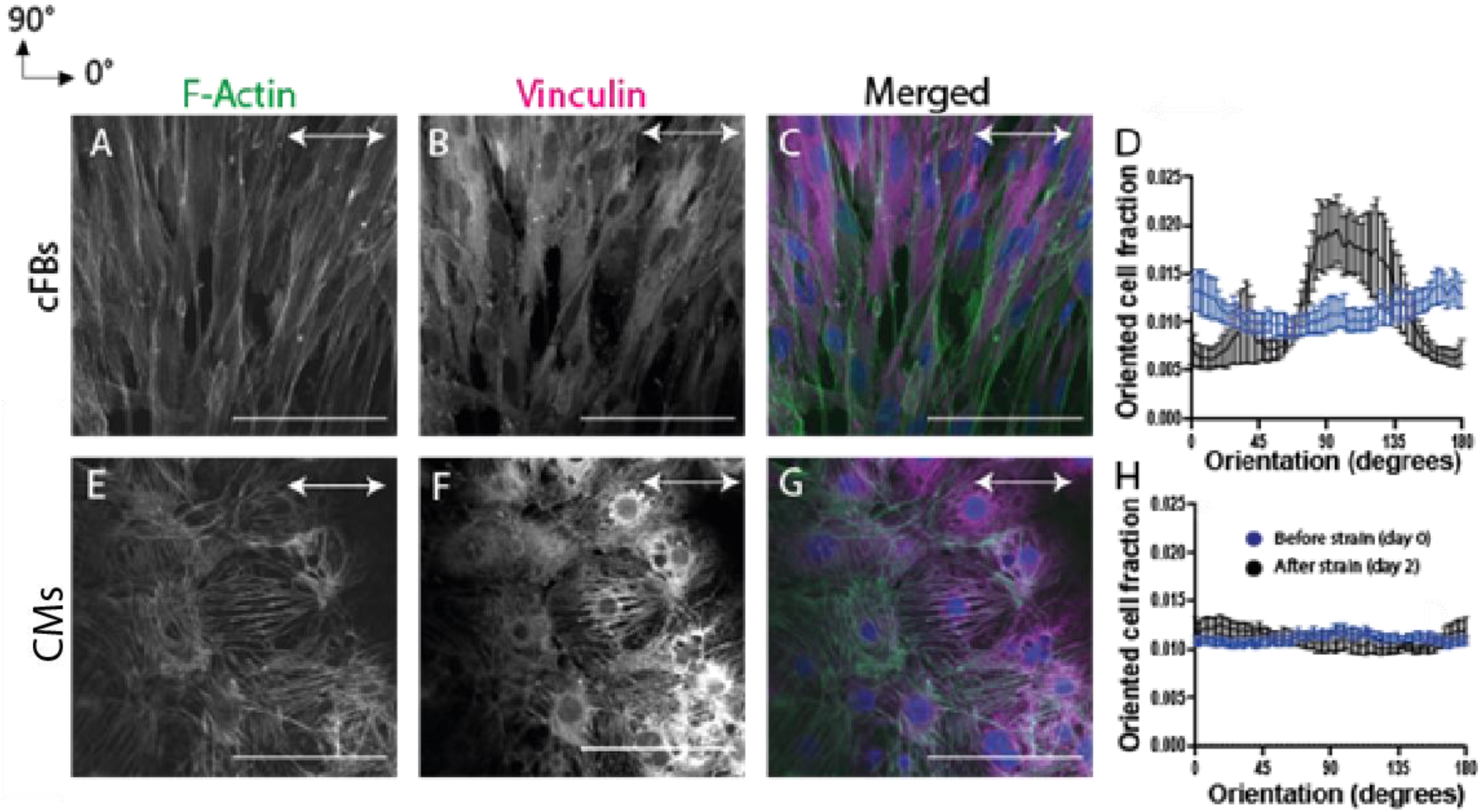
Replication of the experiments with CMs derived from the DRRAGN cell line validated the lack of strain avoidance in CM monocultures upon cyclic strain administration. Representative fluorescent images of the actin cytoskeleton (F-actin) and cell-substrate interactions (vinculin) in strained cFBs (A-C) and CMs (E-G). D, H) Frequency distribution histograms of cFBs (D) and CMs (H) orientation before (blue) and after (black) 48 hours of uniaxial cyclic strain. The direction of the uniaxial cyclic strain is depicted with the white arrow. Scale bar indicates 200 μm. Results are expressed as mean ± SEM (n = 9 from 3 independent experiments).

**Figure S5:**
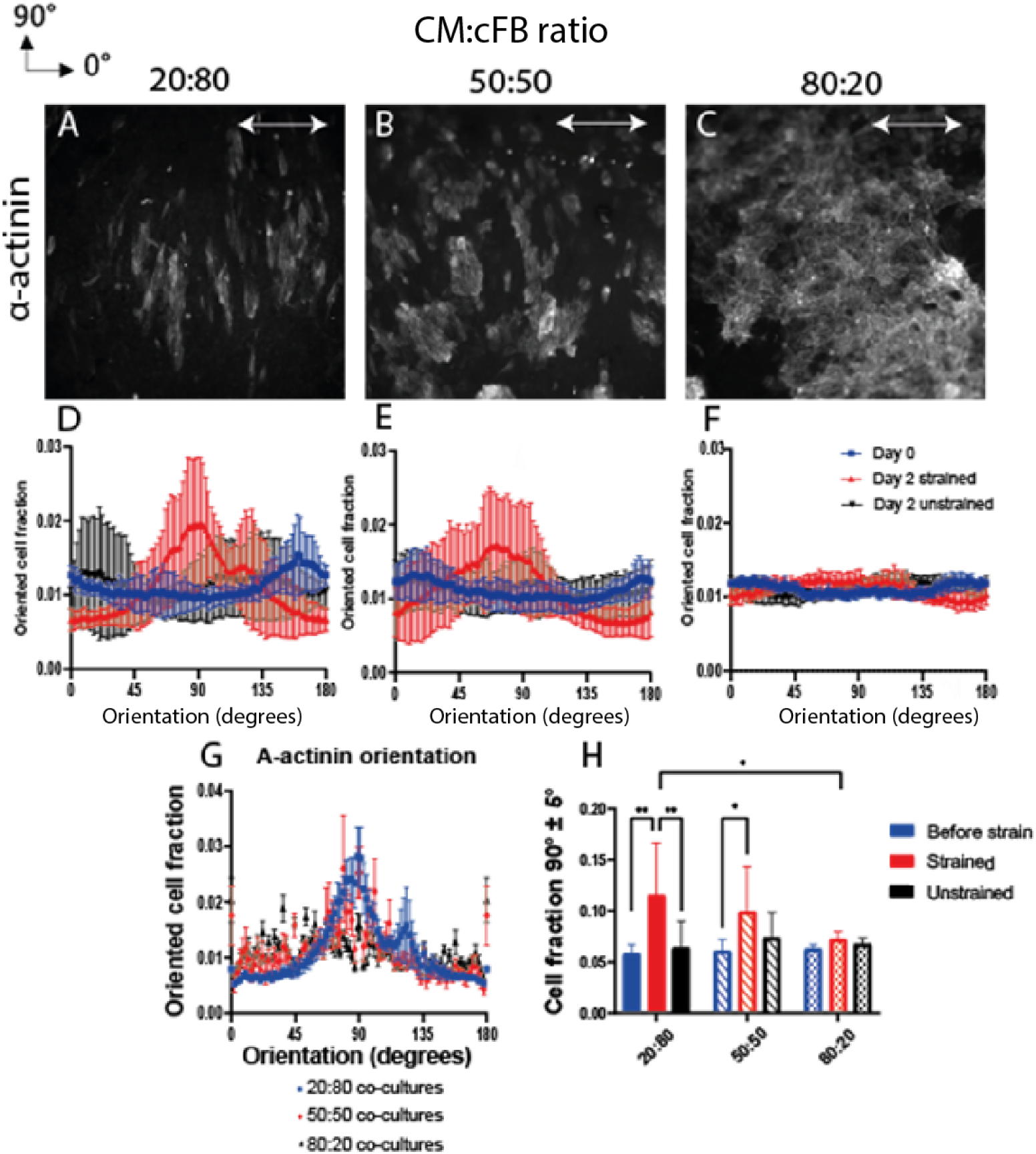
The effect of uniaxial cyclic strain on the organization of co-cultures consisting of 20:80, 50:50, and 80:20 CMs:cFBs ratio. A-C) A-actinin expression in real-time shows the organization of CMs, either as single cell or in clusters, in the 20:80 and 50:50 co-cultures, whereas this is not observed for the 80:20 co-culture. The direction of the uniaxial cyclic strain is depicted with the white arrow. Scale bar indicates 200 μm. D-F) Frequency distribution histograms of the cardiac co-cultures before strain was applied (day 0, blue), after 48 hours of uniaxial cyclic strain (red), and after 48 hours of static culture (black). G) Frequency distribution histogram of α-actinin orientation in strained co-cultures. H) The oriented cell fractions at 90° ± 5° show a significant increase upon 48 hours of uniaxial cyclic strain for the 20:80 and 50:50 co-cultures. Results are expressed as mean ± SEM (n = 6 from 2 independent samples).

**Figure S6:**
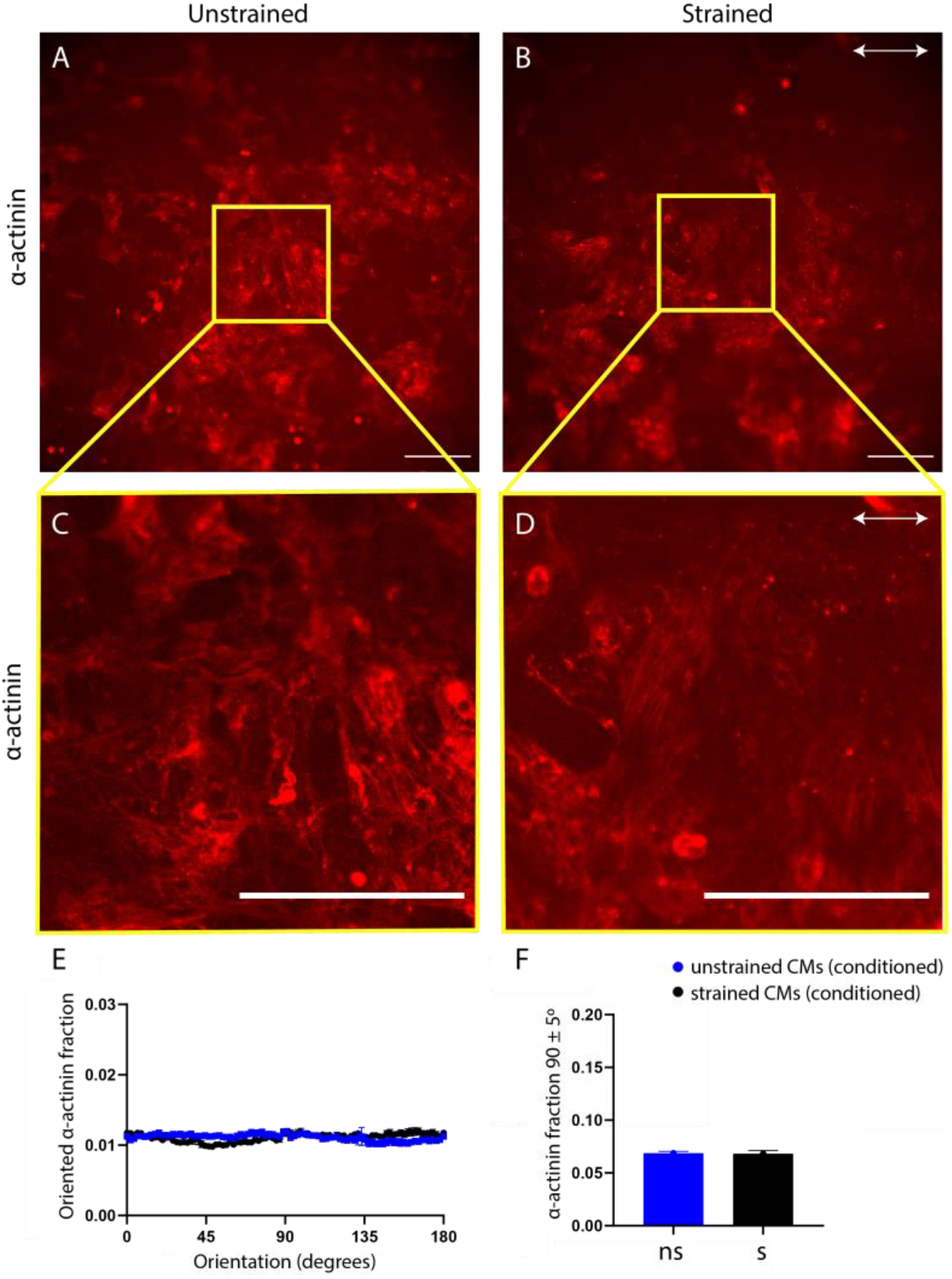
The effect of cFB conditioned medium on the orientation of CMs after 48 hours of static or dynamic culture. A-B) Representative fluorescent images of unstrained (A) and strained (B) CMs (α-actinin, red) that are cultured in cFB conditioned medium, showing randomly oriented CMs in both conditions. E, F) Orientation distribution histogram and oriented cell fraction at 90 ± 5° show no difference in cellular orientation between strained and unstrained CMs cultured in cFB-conditioned medium. N=9 from three independent experiments. Scale bar = 200 μm.

**Figure S7:**
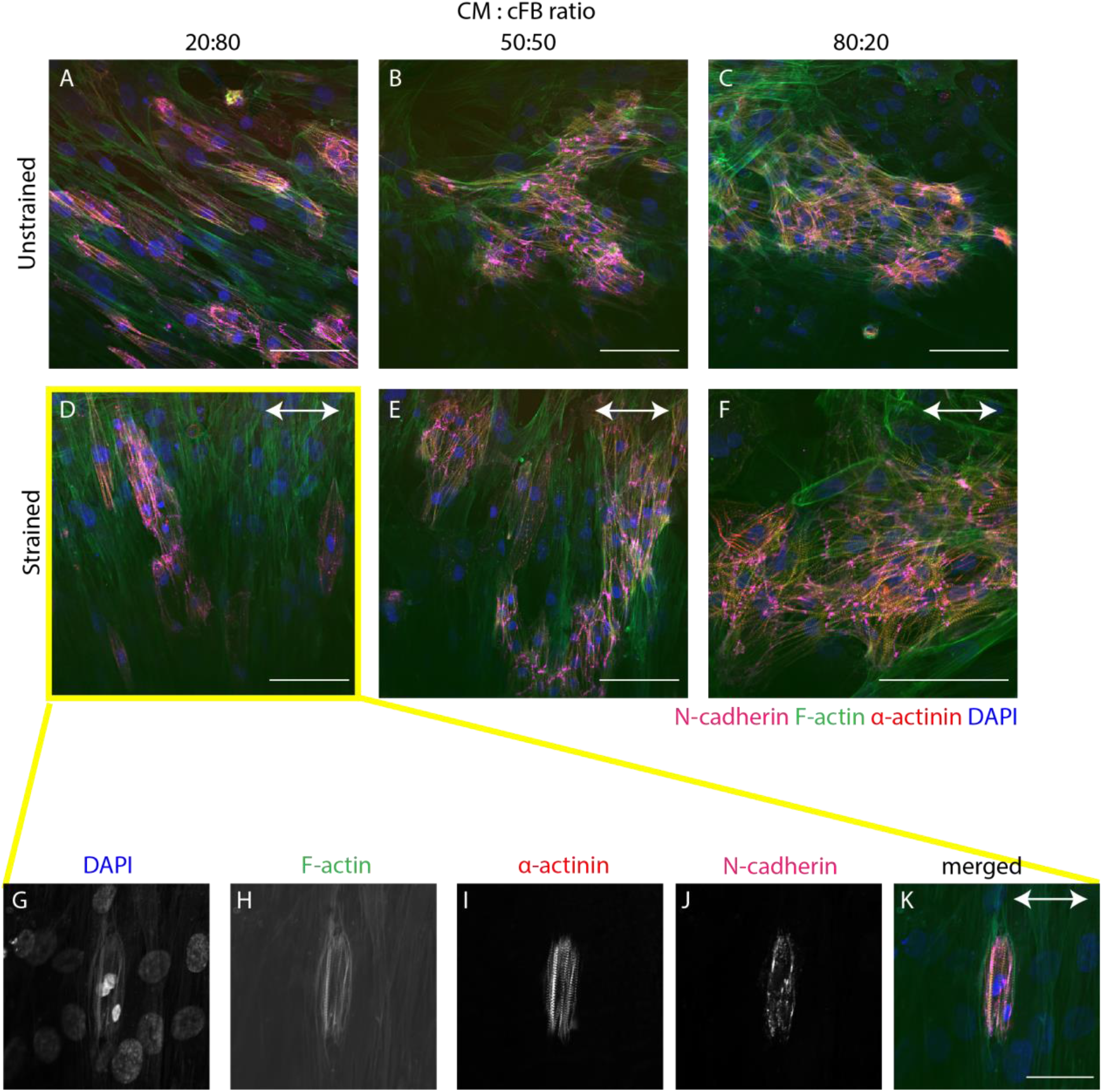
N-cadherin expression in strained co-cultures of varying seeding ratios shows a lack of N-cadherin mediated cell-cell interactions between CMs and cFBs. A-F) Representative fluorescent images of unstrained (A-C) and strained (D-F) co-cultures showing clear N-cadherin expression at the cell membranes solely in CMs concurrent with homocellular cell-cell interactions, which is lacking between cFBs or cFBs and CMs. D-F) Representative fluorescent merged images including F-actin (green) to visualize the cFBs show no N-cadherin expression in cFBs nor at the boundaries between the cFBs and hPSC-CMs. G-K) Isolated CMs surrounded by cFBs lack the typical N-cadherin expression we observe between CMs.

**Figure S8:**
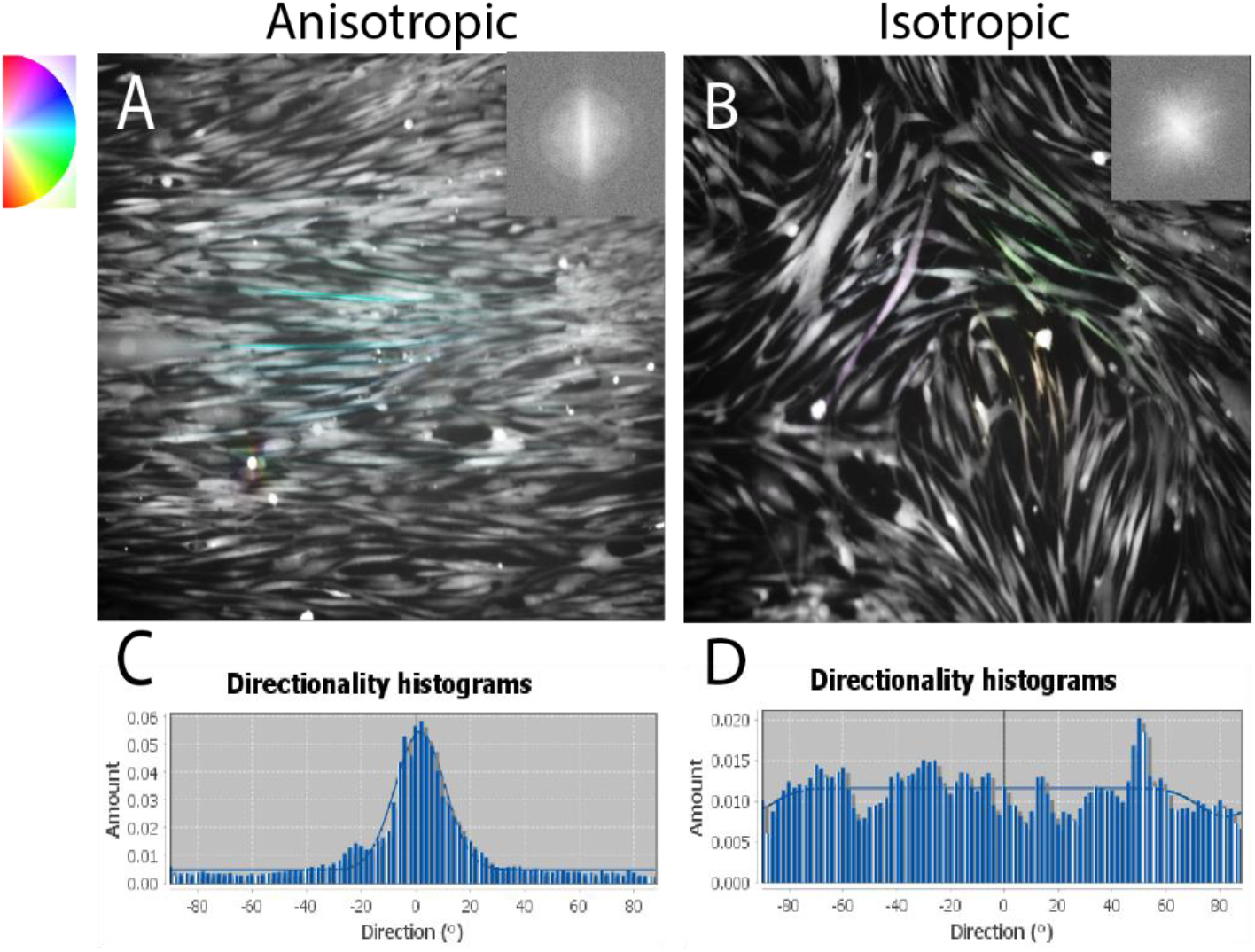
Representative fluorescent images and directionality histograms of linear and randomly oriented cFB to validate directionality quantification method. A – B) Orientation maps of cFBs cultured on either parallel or crosshatch protein patterns are colored according to the local directionality, or location orientation. C – D) Directionality histograms demonstrating either a distinct peak or approaching a flat line for cells cultured on parallel and crosshatch protein patterns, respectively.

**Table S1:** List of all used primary- and secondary antibodies and dyes

| Antigen | Source | Cat.No | Isotype | Label | Species | Dilution |
| --- | --- | --- | --- | --- | --- | --- |
| Vimentin | Abcam | Ab20346 | IgM | - | Mouse | 1/200 |
| A-actinin | Sigma-Aldrich | A7811 | IgG1 | - | Mouse | 1/200 |
| N-cadherin | Sigma-Aldrich | C3865 | IgG1 | - | Mouse | 1/200 |
| Vinculin | Sigma-Aldrich | V9131 | IgG1 | - | Mouse | 1/200 |
| Mouse IgM | Molecular Probes | A21042 | - | Alexa Fluor 488 | Goat | 1/200 |
| Mouse IgG1 | Molecular Probes | A21240 | - | Alexa Fluor 647 | Goat | 1/200 |
| Phalloidin | Thermo-Fisher | - | - | Alexa Fluor 488 | - | 1/400 |
| Phalloidin | Thermo-Fisher | - | - | Alexa Fluor 647 | - | 1/400 |
| DAPI | Sigma-Aldrich | - | - | - | - | 1/500 |

